# The role of neuropeptides in regulating ecdysis and reproduction in the hemimetabolous insect *Rhodnius prolixus*

**DOI:** 10.1101/2022.04.13.488237

**Authors:** Marcos Sterkel, Mariano Volonté, Maximiliano G. Albornoz, Juan Pedro Wulff, Mariana del Huerto Sánchez, Paula María Terán, María Teresa Ajmat, Sheila Ons

**Affiliations:** Laboratorio de Neurobiología de Insectos (LNI), Centro Regional de Estudios Genómicos, Facultad de Ciencias Exactas, Universidad Nacional de La Plata, CENEXA, CONICET, La Plata, Buenos Aires, Argentina; Instituto Superior de Investigaciones Biológicas (INSIBIO). Universidad Nacional de Tucumán. Chacabuco 461, T4000, S. M. de Tucumán, Tucumán

**Keywords:** corazonin, eclosion hormone, ecdysis triggering hormone, orcokinin, crustacean cardioactive peptide, moulting, egg hatching

## Abstract

In ecdysozoan animals, moulting entails the production of a new exoskeleton and the shedding of the old one during ecdysis. It is induced by a pulse of ecdysone that regulates the expression of different hormonal receptors and activates a peptide-mediated signalling cascade. In Holometabola, the peptidergic cascade regulating ecdysis has been well described. However, very little functional information regarding the neuroendocrine regulation of ecdysis is available for Hemimetabola, which displays an incomplete metamorphosis.

Here, we studied neuropeptides related to ecdysis regulation in the hemimetabolous insect *Rhodnius prolixus*. The RNA interference-mediated reduction of ETH expression in fourth instar nymphs resulted in lethality at the expected time of ecdysis, thereby showing its crucial role in this process. Furthermore, the results revealed the involvement of ETH in the regulation of reproductive fitness. Different from holometabolous, the knockdown of ETH in adult females led to failures in egg hatching without affecting the oviposition. Most of the first instar nymphs hatched from the eggs laid by females injected with dsEH, dsCCAP and dsOKA died at the expected time of ecdysis, indicating the crucial involvement of these genes for post-embryonic development. No phenotypes were observed upon CZ knockdown in nymphs or adult females. The conservation of the role of these neuropeptides in regulating ecdysis and reproduction throughout the class Insecta is discussed.

**Summary statement:** The information provided here is of interest for evolutive studies on the neuroendocrine regulation of ecdysis and reproduction in insects, and the research for new targets to control pest insects.

## Introduction

Insects are a predominant life form on Earth; different species are adapted to every terrestrial ecological niche, from deserts to glaciers (Chown, S. L., and Nicolson, 2004). This spectacular adaptability is due to their particular developmental, physiological and reproductive strategies. The exoskeleton, for example, is an excellent adaptation to protect insects from desiccation and physical damage but imposes constrictions for post-embryonic development. To grow and develop insects must shed the old cuticle and emerge as the following instar during a process called ecdysis (Zitnan and Adams, 2012). Moulting involves the synthesis of a new cuticle underneath the old one, the digestion and resorption of most of the old cuticle during pre-ecdysis, and the shedding of the old cuticle during ecdysis. Finally, during post-ecdysis, the cuticle of recently emerged insects becomes dark and rigid (sclerotized) due to melanin deposition and protein cross-linking. Failure to complete ecdysis will be deleterious for the individual, making the regulation of this process an excellent target in the search for new insect pest management strategies (Ewer, 2005; White and Ewer, 2014; Žitňan et al., 2003; Žitňan et al., 2007). Ecdysis is tightly regulated by the neuroendocrine system, i.e., neuropeptides and their receptors. In holometabolous, with minor variations between species, the core components of this neuropeptidergic network are corazonin (CZ), eclosion hormone (EH), ecdysis triggering hormone (ETH) and crustacean cardioactive peptides (CCAP) (Zitnan and Adams, 2012). The accepted model proposes that a peak of ecdysone concentration in haemolymph controls the expression and release of neuropeptides that regulate ecdysis (Kim et al., 2004a; Kim et al., 2006a; Kim et al., 2006b; Kingan et al., 1997; Kruger et al., 2015; Mena et al., 2016; Zitnan et al., 1999; Žitňan et al., 2003). In Lepidoptera, CZ initiates this process by stimulating the release of ETH from Inka cells, whereas, in *Drosophila melanogaster* and *Tribolium castaneum*, this release is independent of CZ (Arakane et al., 2008; Zitnan and Adams, 2012). Increasing concentrations of ecdysone result in the production of ETH in epitracheal Inka cells but prevent its release by suppressing Inka cells’ secretory competence. ETH release is thereby blocked until ecdysone levels decline; there exists an inverse correlation between levels of circulating ecdysone and ETH. High ecdysone levels also sensitize the central nervous system (CNS) to ETH by inducing the expression of the ETH receptors (ETHRs). At the end of pre-ecdysis, ETH acts on target cells, resulting in the release of EH. Via a positive feedback loop between EH and ETH, there is a massive release of both neuropeptides in the haemolymph, which leads to the activation of the neuronal network producing CCAP and the peptide hormone bursicon, which control ecdysis and post-ecdysis behavioural sequences (Fuse and Truman, 2002; Gammie and Truman, 1997; Zitnan and Adams, 2000).

The described model of ecdysis regulation is based on studies performed in Holometabola, mainly *D. melanogaster* (Diptera), *T. castaneum* (Coleoptera), *Manduca sexta* and *Bombyx mori* (Lepidoptera) (Zitnan and Adams, 2012). Even though Holo and Hemimetabola differ in the post-embryonic development, and the latter group comprises numerous species of economic and sanitary relevance, ecdysis regulation has been scattered studied in Hemimetabola. A few studies indicated that ETH and CCAP are involved in ecdysis regulation in hemimetabolous insects (Lee et al., 2013; Lenaerts et al., 2017; Shi et al., 2022; Verbakel et al., 2021), suggesting a conserved regulatory network. Conversely, Orcokinin isoform A (OKA) neuropeptides play a central role in the regulation of ecdysis in *Rhodnius prolixus* (Wulff et al., 2017, 2018) but not in *D. melanogaster* (Silva et al., 2021), pointing to differences in ecdysis regulation between Holo and Hemimetabola.

The expression of neuropeptides involved in ecdysis regulation persists in the adult stage when this process does not still occur. The latter suggests an involvement of these neuropeptides in processes like reproduction. In this sense, a role in ovary maturation has been demonstrated for ETH in *Aedes aegypti* (Areiza et al., 2014), *D. melanogaster* (Meiselman et al., 2017) and *Bactrocera dorsalis* (Shi et al., 2019). Furthermore, ETH receptors (ETHR) are involved in male courtship behaviour in *D. melanogaster* (Deshpande et al., 2019).

*R. prolixus* possess sanitary relevance as a vector of *Trypanosoma cruzi*, the parasite that causes Chagas disease, a neglected life-threatening disease affecting around 8 million people worldwide (World Health Organization, 2020). In addition, this species provides an excellent model for studying development and reproduction in Hemimetabola (Ons, 2017). In the unfed condition, *R. prolixus* remains in a state of arrested development; gorging a blood meal initiates the physiological and endocrinological events leading to ecdysis in nymphs and reproduction in adults, a predictable number of days later. Thus, these events can be precisely timed during experimental designs and studied accurately. Furthermore, gene silencing through RNA interference (RNAi) is highly effective in this species, long-persistent and vertically transmitted from a treated female to the offspring (Paim et al., 2013; Sterkel and Oliveira, 2017; Sterkel et al., 2019). In addition to its relevance for entomological experimentation, RNAi has been proposed as a tool for new-generation insecticides that would be species-specific and environmentally sustainable (Christiaens et al., 2020; Vogel et al., 2019). In the case of triatomines, next-generation approaches for insect control are urgently needed due to the emergence of high pyrethroid resistance and the absence of sustainable alternatives (Capriotti et al., 2014; Fabro et al., 2012; Sierra et al., 2016).

Given the central role of the neuroendocrine system in regulating vital processes, neuropeptides and their receptors are considered putative targets for RNAi-based insecticides (Liu et al., 2020; Scherkenbeck and Zdobinsky, 2009). Here we used RNAi in *R. prolixus* to study the role of three neuropeptides reported to regulate ecdysis in Holometabola: CZ, ETH and EH. Furthermore, we evaluated the role of these neuropeptides along with CCAP and OKA, both involved in ecdysis regulation in *R. prolixus* (Lee et al., 2013; Wulff et al., 2017), in regulating the reproductive fitness of females.

## 2. Materials and methods

### 2.1. Insects rearing

A colony of *R. prolixus* was maintained in our laboratory in a 12 h light/dark schedule at 28 °C and 50-60% humidity. Once a month, the insects were fed on chickens, which were housed, cared for, fed and handled following resolution 1047/2005 (National Council of Scientific and Technical Research, CONICET) regarding the National Reference Ethical Framework for Biomedical Research with Laboratory, Farm, and Nature Collected Animals. This framework is in concordance with the standard procedures of the Office for Laboratory Animal Welfare, Department of Health and Human Services, NIH, and the recommendations established by the 2010/63/EU Directive of the European Parliament related to the protection of animals used for scientific purposes. Biosecurity considerations are in agreement with CONICET resolution 1619/2008, which is in concordance with the WHO Biosecurity Handbook (ISBN 92 4 354 6503).

### 2.2. RNA isolation and cDNA synthesis

Total RNA was extracted from first and fourth instar nymphs (whole-body) using TRIzol reagent (Ambion) according to the manufacturer’s instructions. Following treatment with DNAse I (Promega), first-strand cDNA was synthesized using 1 µg total RNA with MMLV Reverse Transcription Kit (Promega) and poly-T primer, according to the manufacturer’s instructions.

### 2.3. Quantitative PCR

Forward and reverse quantitative PCR (qPCR) primers were designed in different exons to prevent the amplification of genomic DNA that could be present in the sample despite the treatment with DNAseI, except for the *eh* gene, which is composed of one exon. qPCR primers were also designed to amplify a different region of the gene than the primers used for dsRNA synthesis since the injected dsRNA could be retrotranscribed into cDNA and introduce errors in endogenous mRNA quantification. The only exception was the *eh* gene since it was not possible to design efficient primers outside the region amplified for dsRNA synthesis. All primer pairs used for qPCRs were tested for dimerization, efficiency, and amplification of a single product. Only primer pairs with efficiency between 80% and 120 % were used (Table 1). cDNA levels were quantified using FastStart SYBR Green Master Mix (Roche) in an Agilent AriaMx Real-time PCR instrument (Applied Biosystems). The schedule used for the amplifying reaction was as follows: (i) 95°C for 3 min; (ii) 95°C for 15 sec; (iii) 56°C for 15 sec; (iv) 72°C for 30 sec. Steps (ii), (iii) and (iv) were repeated for 40 cycles. Controls without a template were included in all batches. *Elongation factor 1 (EF-1)* and *tubulin (Tub)* were used as housekeeping genes since they were previously validated as stable genes in *R. prolixus* (Majerowicz et al., 2011; Omondi et al., 2015; Paim et al., 2012). The 2e^-ΔCT^ values obtained for treated and control groups were used to evaluate the relative levels of target mRNAs (Livak and Schmittgen, 2001).

**Table 1.**
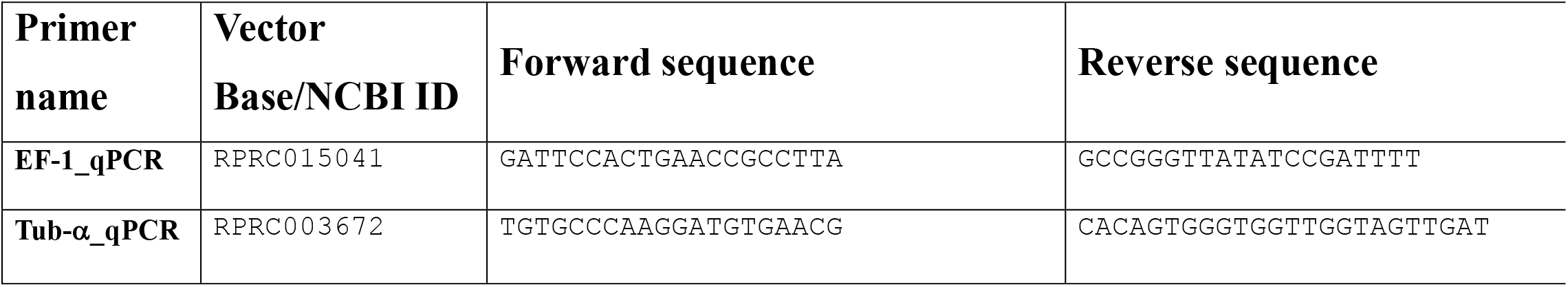
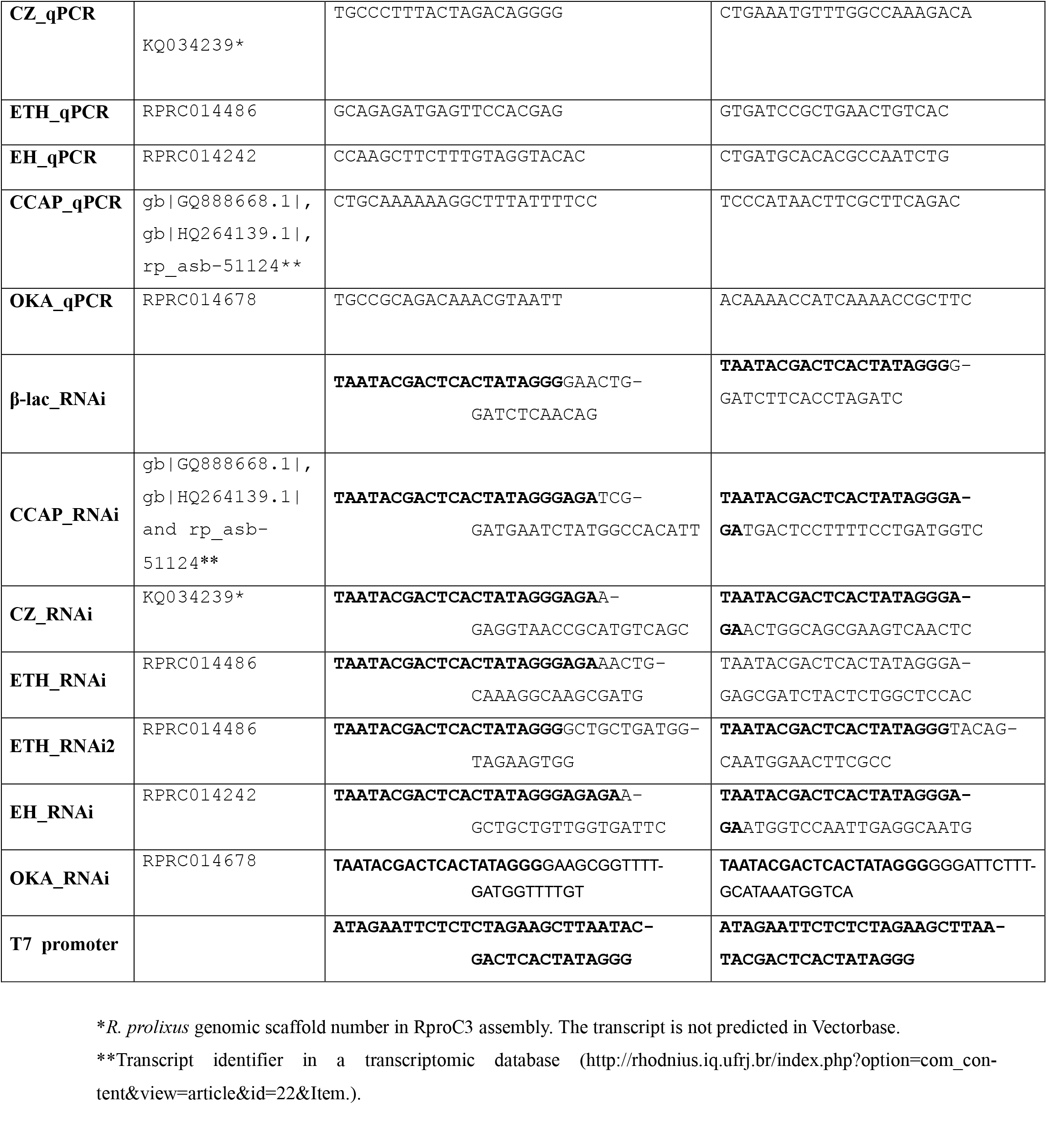
Sequences of primers used in this study. All sequences ID numbers used were as found in Vectorbase database (https://www.vectorbase.org/) or NCBI database (https://www.ncbi.nlm.nih.gov/).

### 2.4 Synthesis of double-stranded RNA (dsRNA)

Specific primers for each target gen were designed (Table 1). The primers for RNAi experiments were designed to knock down all isoforms of each gene, given that three isoforms for *ccap* and two isoforms for *cz* genes have been described (Ons et al., 2011). Three isoforms were described for the *ok* gene that encodes different families of peptides (Sterkel et al., 2012), but RNAi primers were designed to knock down only isoform A (OKA) since only these peptides affect ecdysis in R. *prolixus* (Wulff et al., 2017). A fragment from the *β-lactamase* gene (*β-lac)*, absent in the *R. prolixus* genome, was PCR-amplified from the pBluescript plasmid and used as a control to assess the putative unspecific effects of dsRNA injections. The designed primers contained the T7 polymerase binding sequence at the 5’ end, required for dsRNA synthesis. All PCR products were sequenced (Macrogen, Seoul, Korea) to confirm their identity. One µl of the PCR product was used for a secondary PCR using a T7-full promoter primer (Table 1). The dsRNAs were *in vitro* transcribed using a T7-RNA polymerase (Promega, USA) according to the manufacturer’s instructions. dsRNAs were precipitated with isopropanol and resuspended in ultrapure water (Milli-Q). The dsRNAs were visualized by agarose gel (1.5% w/v) electrophoresis to verify size, integrity and purity. The dsRNAs were quantified from images of the gels using the software ImageJ (National Institute of Health, USA). The dsRNAs were stored at -20°C until use.

### 2.5 RNAi

Fourth instar nymphs and adult females *R. prolixus* were injected into the thorax using a 10 μl Hamilton microsyringe with 2 μg of dsRNA dissolved in 2 μl of ultrapure water. Control insects were injected with 2 μg of β-lac (unspecific) dsRNA. Only mated females that had been fed once during the adult stage were used. Females were injected with dsRNA 21 days after the first blood meal. Fourth instar nymphs and adult females were fed on chicken seven days after the dsRNA injection. The feeding day was considered day cero post-blood meal (PBM). Fourth instar nymphs were collected in Trizol reagent (Ambion, USA) for RNA extraction to check the levels of gene knockdown by qPCR on day 11 PBM, which is the day before the expected moulting in our conditions. A fraction of the first instar nymphs hatched from the eggs laid by dsRNA-injected females were fed on chickens seven days after hatching. On that day, starved first instar nymphs from different groups were collected in Trizol reagent (Ambion, USA) for RNA extraction to check the levels of gene knockdown by qPCR.

### 2.6 Microscopy and histology

Fourth instar nymphs from dsß-lac and dsETH-injected groups were collected on day 12 PBM in Bouin solution (saturated picric acid 70%, formol 25% and acetic acid 5%). Insects from dsß-lac had already moulted to the next instar. After 24 hours, insects were washed three times with ethanol 70%. After washing, insects were kept in ethanol 70% for 6 hours. Insects were transferred to ethanol 90% for 12 hours, ethanol 96% for 2 hours and ethanol 96%: n-butyl alcohol (1:1) for 2 hours. Finally, insects were transferred to n-butyl alcohol for 24 hours. N-butyl alcohol was changed three times every 24 h. Insects were kept in n-butyl alcohol until the microscopies were performed.

For microscopic studies, the samples were dehydrated with ethyl alcohol of increasing strength and butyl acetate was used as an intermediate. A synthetic medium (Histoplast) was used for inclusion, and 4-5 micrometre cuts were made using an SLEE Mainz CUT 4062 microtome. Histological sections were stained with Hematoxylin-Eosin (H-E) and Toluidine Blue. Microphotographs were taken with a Zeiss Primo Star microscope coupled to a Canon Power Shot G10 digital camera.

### 2.7 Survival, ecdysis, oviposition and eclosion measurements

Fully engorged females were individually isolated into vials and kept at 28 °C and 50–60 % relative humidity under a photoperiod of 12 h of light/12 h of dark. The survival and number of eggs laid by each female were recorded daily. The eclosion ratios were calculated as the number of hatched first instar nymphs/number of eggs laid by the female. Nymph survival and ecdysis were also scored daily.

### 2.8 Statistical Analysis

At least two independent experiments were performed for each treatment. Each replica included N=5–15 insects per experimental group. The data from different replicates were combined into a single graph for the design of the figures. The statistics were performed on both, each independent experimental replicate and the final data containing the information from the different replicates combined. The p values indicated in the text are from the data of the independent experiments combined. Statistical analysis and graphs designs were performed using Prism 8.0 software (GraphPad Software). A one-way ANOVA test was used to evaluate differences in mRNA levels during the moulting cycle in fourth instar nymphs. The log-rank (Kaplan-Meier) test was used to evaluate differences in survival rate. A two-way ANOVA test was used to evaluate differences in daily rates of oviposition, eclosion of eggs and ecdysis. An unpaired t-test was performed to evaluate differences between the experimental and control groups in the relative mRNA levels. p≤ 0.05 was considered statistically significant.

## 3 Results

### 3.1 Relative expression of ecdysis-related neuropeptide precursor genes during the moulting cycle in *R. prolixus*

The levels of CZ, ETH and EH mRNAs were measured in 4^th^ instar nymphs at different time points during the moulting cycle: days 4, 6, 8, 10 and 12 PBM. Transcription of the three neuropeptide precursors could be detected throughout the days studied. All the transcripts were significantly upregulated on day 12 PBM, just before the expected ecdysis (n=7) (Figure 1). Besides, ETH and EH mRNA levels were also upregulated on day 10 PBM (n=5) and slightly on day 6 PBM (n=5), in coincidence with the ecdysone peak in haemolymph (Wulff et al., 2017). The augment in mRNA levels during day 6 was more evident for the ETH precursor. This neuropeptide presented higher mRNA levels than CZ and EH in all the time points evaluated (Figure 1).

**Figure 1:**
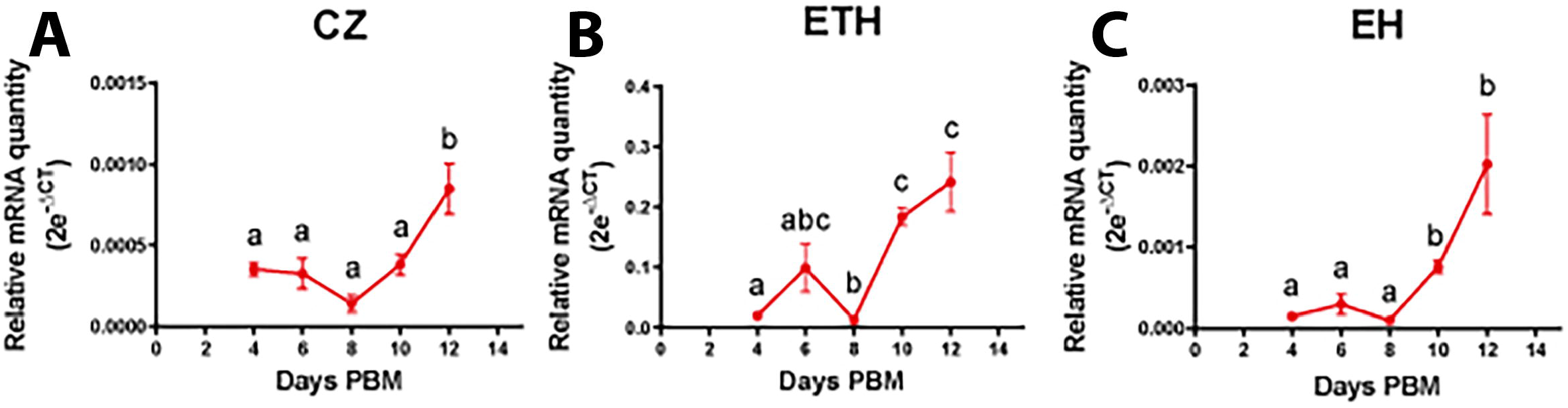
CZ, EH and ETH mRNA levels in fourth instar nymphs during the moulting cycle. The data from multiple experiments were combined into a single graph and plotted as the mean+s.e.m. *Elongation Factor 1 (EF-1) and Tubulin (Tub)* were used as housekeeping genes. Different letters indicate significant differences (one-way ANOVA and DMS contrasts: p<0.05; N= 5-7 in each time point).

### 3.2 Disruption of ETH signalling inhibits ecdysis

Compared to controls on day 11 PBM, dsRNA treatments significantly reduced the transcript levels of ETH (96.44 %. p<0.0001, n=10), CZ (86.44% p<0.0001, n=6) and EH (69.12%. p=0.0448, n=6-11) in 4^th^ instar nymphs (Figure S1). None of the experimental groups showed phenotypic differences with controls until the expected ecdysis period (Figure 2). At that time (day 14 +/-4), most of the insects belonging to the control group (n=45) moulted to normal 5^th^ instar, as did dsCZ (n=14) and dsEH-injected insects (n=25) (Figure 2A). Differently, those insects treated with dsETH (n=61) did not moult. Instead, at the expected time of ecdysis, they stopped any movement and died 1-2 days later (Figure 2A-B). Even though ETH-silenced individuals did not shed the 4^th^ instar cuticle, they presented the 5^th^ instar cuticle hardened and tanned, as revealed by gently removing the upper cuticle with entomological forceps (Figure S2). Histological sections confirmed that apolysis and the partial degradation of cuticle occurred in the arrested specimens, pointing to a defect in the ecdysis itself but not in other moulting-related events (Figure 2 C-D).

**Figure 2:**
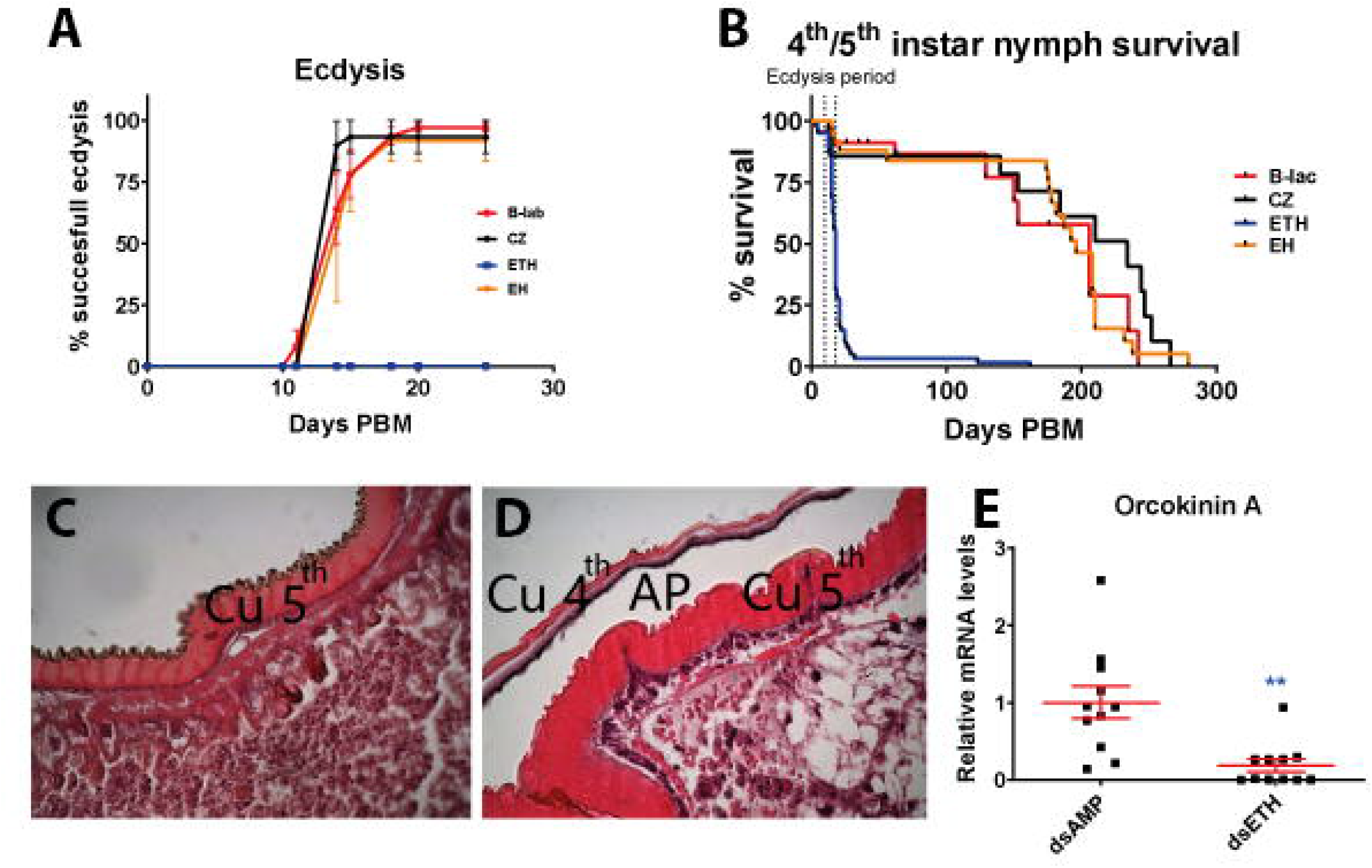
Effects of CZ, ETH and EH knockdown in fourth instar nymphs. **A:** Percentage of successful ecdysis on different days PBM. **B:** Fourth/fifth instar nymph survival. Dotted vertical lines indicate the period when ecdysis occurred in controls. **C:** Cross-section of the cuticle from just emerged control 5^th^ instar **D:** Cross-section of the cuticle from dsETH-injected 4^th^ instar nymph during the arrested ecdysis. Magnification 400x. AP: apolysis space; Cu 4^th^: old cuticle from the 4^th^ instar individual; Cu 5^th^: new cuticle from the 5^th^ instar individual; **E:** OKA mRNA levels on control and dsETH-treated fourth instar nymph 11 days PBM assessed by qPCR. *Elongation Factor 1 (EF-1)* and *Tubulin (Tub)* were used as housekeeping genes. All the 2e^-ΔCT^ values were divided by the average of the 2e^-ΔCT^ values in the control group. ** p<0.01. The data were plotted as the mean+/-s.e.m.

Since the phenotype observed upon dsETH treatment was similar to that previously observed upon the silencing of OKA (Wulff et al., 2017), we measured the mRNA levels of OKA transcript in dsETH-treated and control insects on day 11 PBM. We observed a significant reduction in the expression of OKA in insects belonging to the dsETH group (Figure 2E; p<0.001; n=11). We ruled out off-target effects by injecting dsRNA covering different regions of the ETH transcript, obtaining similar results both in the lethality and phenotype during ecdysis and in the downregulation of the OKA transcript.

### 3.3 Disruption of ETH signalling in females impairs egg hatching

Given that ETH is a gonadotropic factor in the ovaries of holometabolous species (Areiza et al., 2014; M. Meiselman et al., 2017; Shi et al., 2019), we studied the effects of ETH knockdown on the reproductive fitness of *R. prolixus* females. Furthermore, we evaluated the role in *R. prolixus* reproductive fitness of other neuropeptides involved in regulating the ecdysis process in Holo and/or Hemimetabola (CZ, EH, CCAP, OKA). Female survival was not affected in dsETH (n=25), dsCZ (n=13), dsCCAP (n=17) or dsOKA (n=7) groups; a small but significant increase in survival (resistance to starvation. p=0.044, n=15) was observed in females injected with dsEH in comparison with the control group (n=38) (Figure 3 A-B). The egg production was not affected in any experimental group (Figure 3 C-D). However, the egg hatching was dramatically reduced in dsETH-treated animals (p<0.0001, n=32) but not in the dsOKA, dsEH, dsCCAP or dsCZ groups (Figure 3 E-F). We carefully opened several of the unhatched eggs laid by dsETH-injected females to evaluate if they were embryonated or possible defects in the development of the embryos. The eggs were opened on day 12 after being laid (a day when control nymphs had hatched). Interestingly, we observed completely formed pro-nymphs without any evident defect in development (Figure S3), indicating that the impairment in egg hatching was not due to problems in the fertilization or the embryonic development. Off-target effects were ruled out by injecting dsRNA covering different regions of the ETH mRNA, achieving identical phenotypes.

**Figure 3:**
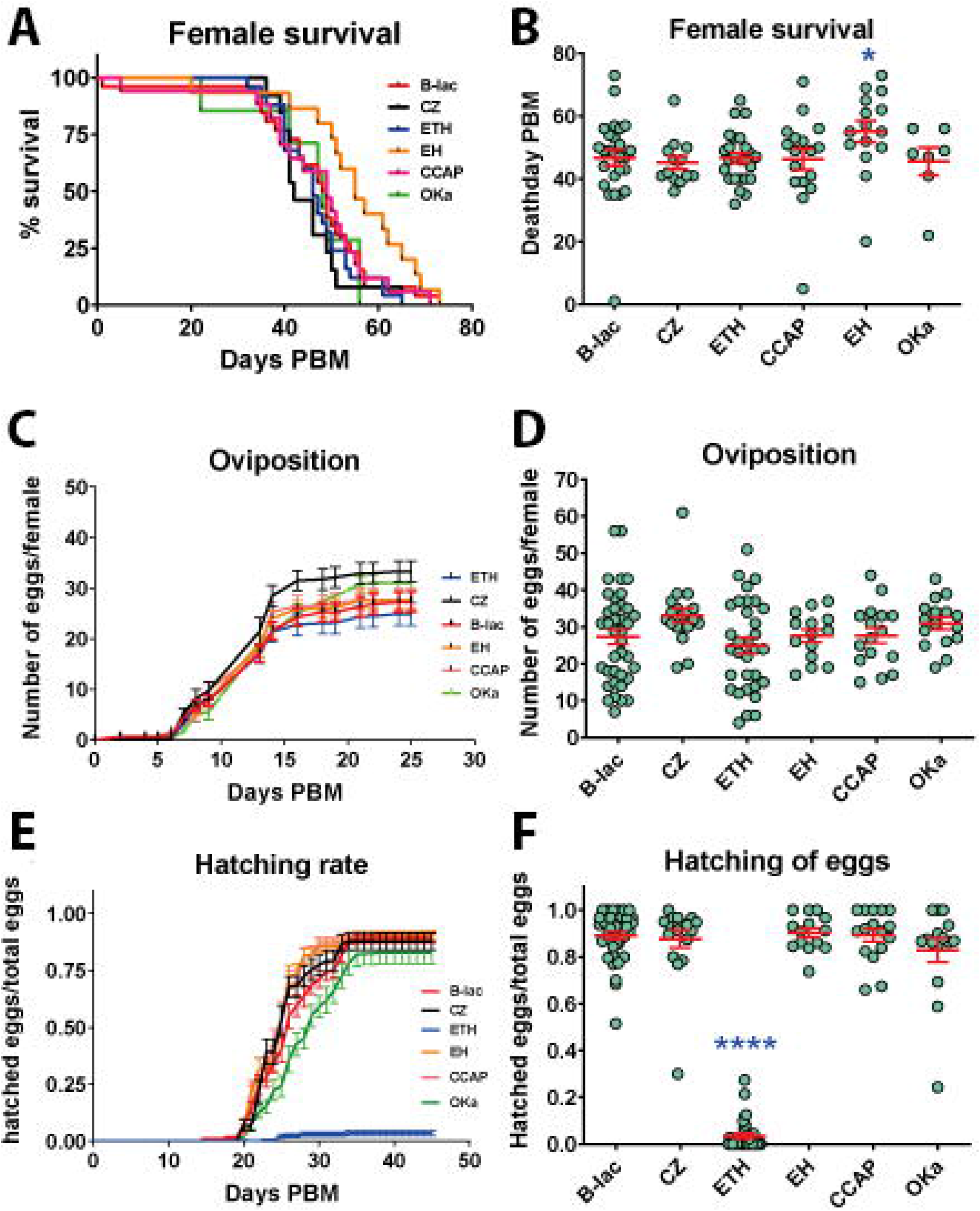
Effects of CZ, ETH, EH, CCAP and Oka knockdown in females. **A:** Female survival Kaplan-Meier curves. **B:** Number of days PBM when dead was registered in females injected with different dsRNAs. **C:** Cumulative number of eggs laid daily by *R. prolixus* females over days PBM. **D:** Total number of eggs laid daily by *R. prolixus* females after 25 days PBM. **E:** Proportion of eggs hatched over days PBM. **F:** Proportion of eggs hatched after 25 days PBM. The data were plotted as the mean+s.e.m. * p<0.05. **** p<0.0001.

### 3.4 Effect of vertical transmission of neuropeptides silencing on 1^st^ instar nymphs

qPCR determinations revealed reduced levels of target mRNAs in the 1^st^ instar nymphs hatched from dsRNA-treated females for dsCZ (87.9%), dsEH (46.5%), dsCCAP (97.6%) and OKA (80%) (Figure S4). We fed first instar nymphs and evaluated the effects on survival and ecdysis. Unexpectedly, different from 4^th^ instar nymphs, 83% of the nymphs silenced for EH expression died during the expected ecdysis period without performing ecdysis (n=58; p<0.0001). This phenotype is comparable to the one observed upon ETH knockdown in 4^th^ stage nymphs (Figure 2). Similar results were observed for OKA (88.9% mortality. n=18; p<0.0001) and CCAP-silenced nymphs (86.8% mortality. n=38; p<0.0001. Figure 4A-B). As observed in 4^th^ instar nymphs, most of the control (n=55) and CZ-silenced 1^st^ instar nymphs moulted normally to the 2^nd^ instar, even though the survival of the latter group was reduced (n=38; p=0.0077. Fig. 4A-B). Unfed nymphs (which do not start the moulting process) didn’t present differences in survival rates among the different experimental groups (Figure 4C). This fact also indicates that the reduction in survival rate was due to failures during ecdysis in the blood-fed nymphs. Recording the moulting rate for the ETH group was not possible given the almost null rate of egg hatching.

**Figure 4:**
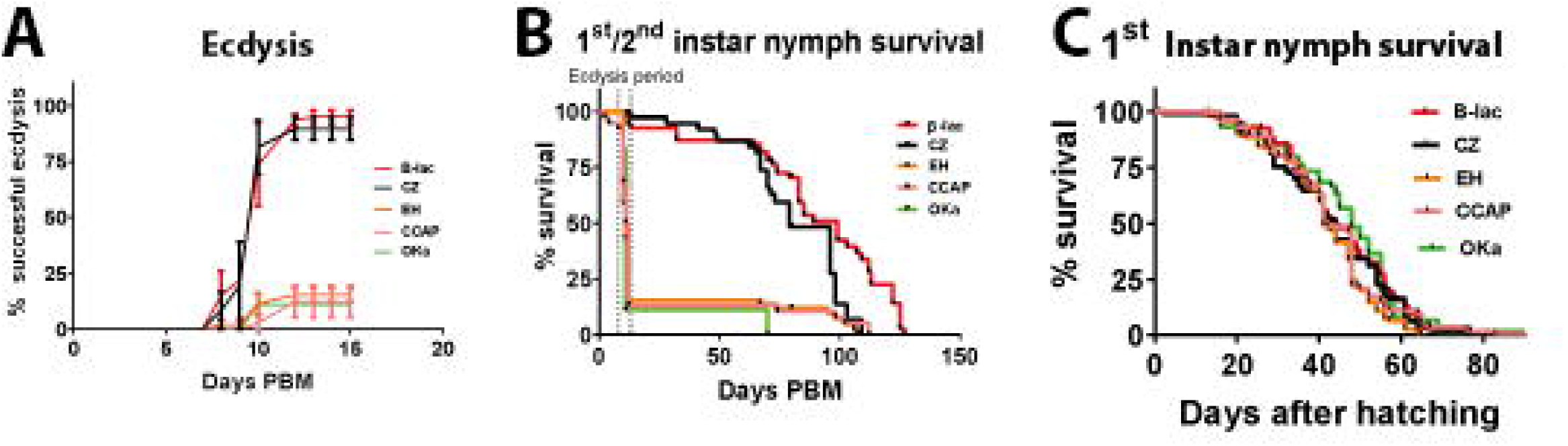
Effects of CZ, EH, CCAP and OKa Knockdown in first instar nymphs. **A:** Percentage of successful ecdysis on different days PBM. **B:** 1^st^/2^nd^ instar nymph survival. Dotted vertical lines indicate the period when ecdysis occurred in controls **C:** unfed 1st instar nymph survival.

## 4. Discussion

Ecdysis and reproduction are critical processes in the life cycle of insects, which are much better studied in holo than in hemimetabolous. *R. prolixus* is a convenient model to study post-embryonic development and reproduction, and it is very sensitive to gene silencing by systemic RNAi (Ons, 2017). These facts allowed us to study the role of different neuropeptide precursors, which were involved in ecdysis in holometabolous insects (Zitnan and Adams, 2012), using an appropriate hemimetabolous model. Even though mRNAs coding for CZ, ETH and EH were detected in every time point analysed throughout the 4^th^ instar moulting cycle, they were significantly upregulated 1-2 days before ecdysis. ETH mRNA levels were also upregulated on day 6 PBM coinciding with the ecdysone peak in haemolymph (Wulff et al., 2017). The correlation between ecdysone peak and upregulation of ETH expression was also observed in Holometabola and *Schistocerca gregaria* (Lenaerts et al., 2017; Zitnan and Adams., 2012). Silencing of ETH or ETHR in different holometabolous insects resulted in lethality at the expected time of ecdysis (Arakane et al., 2008; Diao et al., 2016; Zibaee et al., 2017), as well as in the hemimetabolous *S. gregaria* (Lenaerts et al., 2017). Our results reinforce the evidence indicating that the role of ETH in the control of ecdysis is conserved across the class Insecta. Those specimens that failed to moult exhibited the duplicated cuticle phenotype characteristic of interrupted ecdysis (Wulff et al., 2017). Histological sections showed that they could perform other events of the moulting process (apolysis, digestion of the old cuticle and synthesis of the new one). However, they failed to shed out the exuvia and died.

The similarity in the phenotypes obtained in 4^th^ instar nymphs upon gene-silencing suggested that a functional relation between ETH and OKA could exist. Indeed, we found that dsETH treated insects had reduced levels of OKA transcripts on day 11 PBM. In our previous work, we demonstrated that the silencing of OKA negatively affected the expression of ETH on day six but not on day 11 PBM (Wulff et al., 2018). Altogether, the results could suggest positive feedback between OKA and ETH to act on the regulation of ecdysis in *R. prolixus*. Interestingly, a recent survey demonstrated that the expression profiles of ecdysis-related neuropeptides correlate positively with each other in a wide range of species (Zieger et al., 2021); the results presented here confirm this tendency for *R. prolixus*. Given that the down-regulation of OKA expression doesn’t affect ecdysis in *D. melanogaster* (Silva et al., 2021) nor in *Blattella germanica* (Ons et al., 2015), positive feedback seems to be not conserved among insects. Further experiments with species from different Orders will be interesting to infer the role of OKA signalling throughout the evolution of ecdysis regulation.

A gonadotropic role of ETH has been demonstrated recently for dipteran species (Areiza et al., 2014; M. Meiselman et al., 2017). ETH signalling deficit in *D. melanogaster* reduced egg production and ovary size, caused by a diminution in Juvenil Hormone levels (M. R. Meiselman et al., 2018). In parallel, OKA silencing affects vitellogenesis and ovary maturation in the cockroach *B. germanica* (Ons et al., 2015) and egg production in *D. melanogaster* (Silva et al., 2021). We did not observe differences in egg production either in ETH-silenced insects or in the animals expressing diminished levels of EH, OKA, CZ or CCAP, pointing to different endocrine regulation of ovary maturation in *R. prolixus*. However, the hatching rate was significantly affected in dsETH-treated females, despite full developed nymphs were be observed in the eggs upon manual dissection, indicating that embryo development proceeded correctly until the late stages of embryogenesis. In hemipterans and other hemimetabolous, shedding of the embryonic cuticle occurs during egg hatching (Truman & Riddiford, 2019). This process is called pronymphal ecdysis, which requires a coordinated activity. The role of neuropeptides in regulating this pronymphal ecdysis has not been studied to date, even though a significant upregulation of ecdysis-related neuropeptides occurs during egg hatching in Hexapoda species with direct development (Zieger et al., 2021). Our results indicate that the insects died inside the egg during the pronymphal ecdysis when ETH was silenced. Since this neuropeptide is involved in nymphal ecdysis, our results suggest a similar regulation of pronymphal and nymphal ecdysis.

The role of CZ in insects is less well clarified when compared to ETH. Different reports include the involvement of CZ in the regulation of feeding, nutrient-sensing, nutritional and oxidative stress, ethanol-related behaviour and metabolism, sperm transfer and copulation, and fecundity (reviewed by (Zandawala et al., 2018). In Lepidoptera, CZ initiates the ecdysis sequence by stimulating the release of ETH from Inka cells (Kim et al., 2004b), even though this role was not observed in *D. melanogaster* (Zitnan & Adams, 2012). Given that in *T. castaneum* and other coleopterans CZ hormonal system seems to be absent, it was proposed that the effect of CZ in ecdysis initiation would be specific for Lepidoptera (Arakane et al., 2008). However, a recent report points to the involvement of CZ in larval-pupal transition in *Bactrocera dorsalis* (Diptera: Tephritidae) via the regulation of ETH expression in Inka cells (Hou et al., 2017). This fact suggests that the role of CZ in post-embryonic development could be conserved in different insect groups. The results presented here indicate that CZ function is not required for ecdysis in *R. prolixus*. In agreement, Hamoudi and colls. (Hamoudi et al., 2016) did not find significant effects on ecdysis when the CZ receptor was silenced in the 4^th^ instar nymphs *R. prolixus*.

Injections of EH in lepidopteran species (*M. sexta* and *B. mori*) caused premature ecdysis (Copenhaver & Truman, 1982; Fugo A., 1983), whereas EH silencing in *T. castaneum* suppressed ecdysis (Arakane et al., 2008). These results suggest that EH is necessary and sufficient to induce ecdysis in insects. However, in *D. melanogaster*, EH ablation in Vm neurons caused minor defects in larval ecdysis (Clark, 2004; McNabb et al., 1997). More recently, Krüger et al. (2015) isolated null EH mutants that died at the time of ecdysis. This apparent contradiction was recently solved with the identification of a broader expression of EH, previously supposed to be restricted to Vm neurons (Scott et al., 2020). The authors identified EH expression in neurons other than the Vm at the adult stage and in somatic tissues, mainly tracheae, at all *D. melanogaster* instars. When EH expression was suppressed in larvae 100% of the animals died, many of them with signals of failure in cuticle shedding. By contrast, nearly 50% of the animals were able to moult to the adult stage when EH expression was restricted to the period of pupal development (Scott et al., 2020). In agreement, we obtained 83% of lethality at the expected ecdysis time when EH was silenced in *R. prolixus* during 1^st^ nymphal instar, but not observable phenotype when 4^th^ instar nymphs were injected with dsEH. In agreement, Zieger et. al. (2021) reported a higher expression of ecdysis-related neuropeptides in the early instar of direct development of Hexapoda compared to the late juvenile stage. In conjunction, recent evidence (present results; Scott et al., 2021; Zieger et al., 2021) suggests that younger instar could be more sensitive to EH signalling than later stages.

The accepted model of neuroendocrine regulation of ecdysis in holometabola proposes that the primary regulator of EH secretion is ETH (Zitnan & Adams, 2012). However, recent results revealed EH expressing cells that do not express ETH receptors, suggesting an alternative regulatory mechanism (Scott et al., 2020). The phenotypes we obtained upon ETH and EH silencing were different, indicating that this could also be the case in *R. prolixus*. Finally, in agreement with previous observations in *R. prolixus* 4^th^ instar nymphs (Lee et al., 2013), the silencing of CCAP transcription provoked deficits of ecdysis behaviour in a significant percentage of 1^st^ instar animals.

Despite the enormous differences in post-embryonic development between Holometabola and Hemimetabola, our results suggest high functional conservation in the neuropeptides that regulate ecdysis. The only exception is OKA, which seems to be essential for successful ecdysis in *R. prolixus* (present results, (Wulff et al., 2017, 2018) but not in *D. melanogaster* (Silva et al., 2021). However, the involvement of OKA in ecdysis could be restricted to Hemiptera or Heteroptera, given that this phenotype was not observed in *B. germanica* (Ons et al., 2015). Different from other species (M. Meiselman et al., 2017; M. R. Meiselman et al., 2018; Ons et al., 2015), neither ETH nor OKA silencing affected ovary development or oviposition in *R. prolixus*, but ETH knockdown prevented eggs hatching, probably by affecting pro-nymph ecdysis.

The information provided here is of interest both, for evolutive studies on the regulation of ecdysis in insects and also in the research for new targets for pest insect management strategies that would be safe for vertebrates and other non-ecdysozoans. The phenotypes observed in survival and reproduction upon ETH silencing indicate that it could be an excellent target for the control of triatomine vectors.

## 6. Acknowledgments

We thank Raúl Stariolo and Patricia Lobbia from Centro de Referencia de Vectores for their generous assistance in the rearing of insects.

## 7. Competing interests

The authors declare no competing interests.

## 8. Funding

This work was supported by grants from the Consejo Nacional de Investigaciones Científicas y Técnicas (CONICET Grant PIP 2015 076 to S.O and CONICET Grant PIP 11220200101744CO to M.S), Agencia Nacional de Ciencia y Tecnología (ANPCyT Grant PICT startup 2018-0275 and 2018-0862 to S.O. and ANPCyT Grant PICT 2017-1015 and 2018 04354 to M.S).

## 9. Author Contributions

M.S designed and performed the experiments, analysed the data and wrote the paper. M.V performed the experiments and contributed critical discussion of the experiments and the data. M.G.A performed some RNAi experiments. J. P. W performed and analysed the expression pattern of neuropeptides in 4^th^ instar nymphs. M.H.S, P.M.T and M.T.A carried out the microscopy studies. S.O designed the experiments and wrote the paper. All of the authors discussed the results and read and contributed to the final version of the manuscript.

## Notes

### Competing Interest Statement

The authors have declared no competing interest.

